# Type VIIb secretion system recruits a dedicated cell wall hydrolase EssH to enable effector secretion by *Staphylococcus aureus*

**DOI:** 10.1101/2025.10.30.685510

**Authors:** Richard Agyen, Isabelle Powell, Mahalia McNair, Dominique Missiakas, Maksym Bobrovskyy

## Abstract

*Staphylococcus aureus* is a pervasive human pathogen that heavily relies on protein secretion to exert its virulence strategies. *S. aureus* encodes a specialized type VIIb secretion system (T7SSb) that contributes to virulence and persistence in the host. T7SSb supports secretion of small proteins belonging to a WXG100 family as well as larger LXG polymorphic toxins. Secretion of these proteins is facilitated by the T7SSb complex that assembles in the cell envelope from core components EsaA, EssA, EssB, and the conserved ATPase EssC. T7SSb-mediated secretion also requires the cell wall hydrolase EssH that bears a cystine histidine-dependent amidase/peptidase (CHAP) domain. Hereby, we show that the N-terminal domain of EssH functions together with the CHAP domain to support T7SSb secretion. We find that EssH copurifies with the T7SSb core components and is required for the assembly of EsaA across the cell wall. Finally, secreted EssH is released into the extracellular milieu and degraded by the secreted staphylococcal protease, staphopain A. We propose a model to capture the function of the dedicated cell wall hydrolase EssH for T7SSb in *S. aureus*.

**IMPORTANCE:** *Staphylococcus aureus* is a leading cause of infections worldwide. *S. aureus* utilizes a specialized type VIIb secretion system (T7SSb) to persist in the infected host tissues as well as target competitor bacteria to establish its niche. T7SSb assembles into a multiprotein translocation complex and facilitates secretion of a set of small proteins and larger polymorphic toxins across the cytosolic membrane. Beyond the membrane, secreted proteins were thought to diffuse through the thick yet porous cell wall and release into the environment. Here, we demonstrate for the first time that *S. aureus* T7SSb extends across the cell wall via its EsaA subunit. Furthermore, accommodation of EsaA within the cell wall requires an associated cell wall hydrolase EssH and is essential for protein secretion via T7SSb. Thus, our findings provide a mechanistic insight for a coordinated cell wall processing and T7SSb assembly to support specialized protein secretion in *S. aureus*.

## INTRODUCTION

Protein secretion is an essential process in bacteria, whereby proteins synthesized in the cytosolic compartment are selectively translocated across the membrane barrier. This process commonly involves protein complexes that form channels across lipid bilayer and utilize ATP or the proton motive force to power protein translocation. In general, precursor proteins that possess N-terminal signal peptides are recognized by the SecA ATPase and delivered to the membrane-spanning SecYEG translocon (1). In Gram-negative organisms, proteins engaged by the Sec pathway are secreted into the periplasmic space, the compartment between the plasma and outer membranes. Gram-negative organisms evolved specialized secretion systems classified as Types I-VI, to secrete proteins into the extracellular milieu or into target cells (2). In Gram-positive organisms, precursor proteins with a single N-terminal sequence are also threaded through the SecYEG translocon and released in the extracellular milieu; indeed, most protein transit across the murein sacculus is not thought to require any additional catalyst (3). Still, Gram positive organisms have evolved specialized protein export systems including type VII secretion systems (T7SS). A T7SS was first described in the diderm-mycolate Actinobacteria and later in monoderm Firmicutes, and are now recognized as two structurally distinct subtypes, T7SSa and T7SSb, respectively (4). T7SSs have since emerged as critical virulence determinants in several important pathogens including *Mycobacterium tuberculosis*, *Streptococcus pyogenes* and *Staphylococcus aureus*, where it facilitates the export of diverse effector proteins essential for host-pathogen interactions (5), nutrient acquisition (6), and bacterial competition (7).

*S. aureus* is a human commensal and pathogen associated with severe invasive infections of the skin, bone, soft tissue and bloodstream (8). In *S. aureus* the T7SSb has been named ESAT-6-like secretion system and is organized in a gene cluster that encodes the translocation machinery components and several secreted substrates (9, 10). Mutants of *S. aureus* defective in T7SSb display reduced virulence and elicit altered pro-inflammatory immune responses during bloodstream infection in mice (9, 11). T7SSb effectors have also been implicated in bacterial antagonism, although it is not yet clear how this property contributes to *S. aureus* colonization and pathogenesis (12, 13). Thus far, six T7SSb effectors are described in *S. aureus* USA300, a major methicillin-resistant *S. aureus* (MRSA) lineage associated with community-acquired infections. These effectors belong to two families: 1) WXG100-like substrates EsxA, EsxB, EsxC and EsxD and 2) LXG-like polymorphic toxins EssD (EsaD) and TspA (9–12, 14–17). WXG100-like proteins are small, 10-15 kDa, and were reported to form homodimers (EsxA-EsxA and EsxC-EsxC) and heterodimers (EsxA-EsxC and EsxB-EsxD) (18). While the physiological roles for these proteins are not well defined, streptococcal EsxA has been noted to exhibit membrane pore-forming properties (5) and staphylococcal EsxB targets the STING pathway in infected cells (19). The polymorphic toxins EssD (68 kDa) and TspA (52 kDa) are larger and contain C-terminal toxic domains with nuclease and membrane depolarizing activities, respectively (12, 15, 17). Bioinformatic analysis identified a genetic association of EssD and TspA toxins with their cognate immunity factors EssI (EsaG) and TsaI, which assure protection from self-intoxication (12, 15, 17). Secretion of WXG- and LXG-like proteins relies on the assembly of T7SSb membrane complex. Affinity purification of tagged EssB and EssC proteins from detergent-solubilized *S. aureus* membranes indicates that four transmembrane (TM) proteins EsaA, EssA, EssB and the FtsK/SpoIIIE-like P loop ATPase EssC, as well as EsxA and the cytosolic, ubiquitin-like protein EsaB, associate into a complex (20, 21). Based on structural studies of the EccC homologue in mycobacteria, EssC is thought to form a central hexameric pore in the cytosolic membrane and utilizes ATP to translocate effector proteins across the lipid bilayer (22, 23). Evidence from *Bacillus subtilis* and *Streptococcus gallolyticus* suggest that EsaA may further extend the T7SSb complex beyond the cell wall (24, 25).

In *S. aureus*, the murein sacculus or cell wall is comprised of a highly cross-linked peptidoglycan (PG) substituted with wall teichoic acid (WTA) and Sortase A-anchored surface proteins (26). PG is synthesized by glycosyltransferases, such as bifunctional penicillin binding protein 2 (PBP2) and monofunctional MGT and SgtA, that polymerize lipid II precursor [C_55_-(PO_4_)_2_-MurNAc-(L-Ala-D-iGln-(NH_2_-Gly_5_)L-Lys-D-Ala-D-Ala)-GlcNAc] to form glycan chains of β-(1→4)-linked repeats of N-acetylglucosamine (GlcNAc) and N-acetylmuramic acid (MurNAc) (27–30). Disaccharide pentapeptide chains are cleaved to 6-8 units in length by SagB (31, 32) and cross-linked in a transpeptidation reaction by PBP1, PBP2 and PBP4 that cleave D-Ala-D-Ala amide bonds allowing for the carboxyl group of D-Ala at position four to bond with the amino group of the pentaglycine cross bridges of the adjacent PG strands (33–36). Recently, we identified EssH, a conserved PG hydrolase associated with the ESS locus of *S. aureus* (37). EssH contains a C-terminal <u>c</u>ysteine, <u>h</u>istidine-dependent <u>a</u>midohydrolase/<u>p</u>eptidase (CHAP) domain which cleaves the MurNAc-L-Ala amide bond within the wall peptide as well as peptide bonds after the first and the fourth glycyl residues within the pentaglycine cross bridge (37). Despite its PG hydrolytic activity, EssH is not lytic unlike some phage-encoded endolysins (37). Importantly, *essH* mutants display defects in protein secretion via T7SSb. While EssH is required for T7b-dependent secretion, it is not a T7SSb substrate. Instead, EssH bears a canonical N-terminal signal peptide targeting it for secretion via the general Sec pathway. Hereby, we performed the systematic molecular characterization of EssH to gain insight into how it coordinates its secretion and activity to support T7SSb substrate transport.

## RESULTS

### EssH is secreted under T7 permissive conditions

Using western blotting approaches to identify T7SSb proteins, we noted earlier that immune detection varies depending on growth conditions (37). Such an example is shown in Fig. 1A with a culture of *S. aureus* USA300 LAC* (hereby referred to as wild type) fractionated into cell and medium (*i.e.*, the pellet and spent culture medium). Culturing *S. aureus* in tryptic soy broth supplemented with heat-inactivated horse serum (HS/HI) at pH 5.5 instead of pH 7 allows for the clear immune detection of EsxA in the medium that contains secreted proteins (Fig. 1A). For sake of simplicity, we refer to the two growth conditions as T7 permissive (pH 5.5) and T7 non-permissive (pH 7). Interestingly, the secretion of EssH also follows a similar pattern, and is detected in the medium only under T7 permissive conditions (Figs. 1A and 1B). Further, when using isogenic mutants *ΔesxA-0304* (lacking all the T7SSb components) and *essH*, we confirmed that the secretion of EssH is T7 permissive but T7-machinery independent (Fig. 1A, lanes 2), yet EssH secretion is required for the secretion of EsxA (Fig. 1A, lanes 3). Next, we asked if the T7 permissive medium may simply induce the production of EssH. To examine this possibility, *essH* was expressed under a constitutive promoter and cloned on a plasmid. However, expression of *essH in trans* also failed to restore EssH and EsxA secretion under T7 non-permissive conditions (Fig. 1C). Thus, simply increasing the copy number or transcription of the *essH* gene cannot enhance its own secretion or that of EsxA suggesting that the culture condition does not directly control the expression of *essH*.

**Figure 1.**
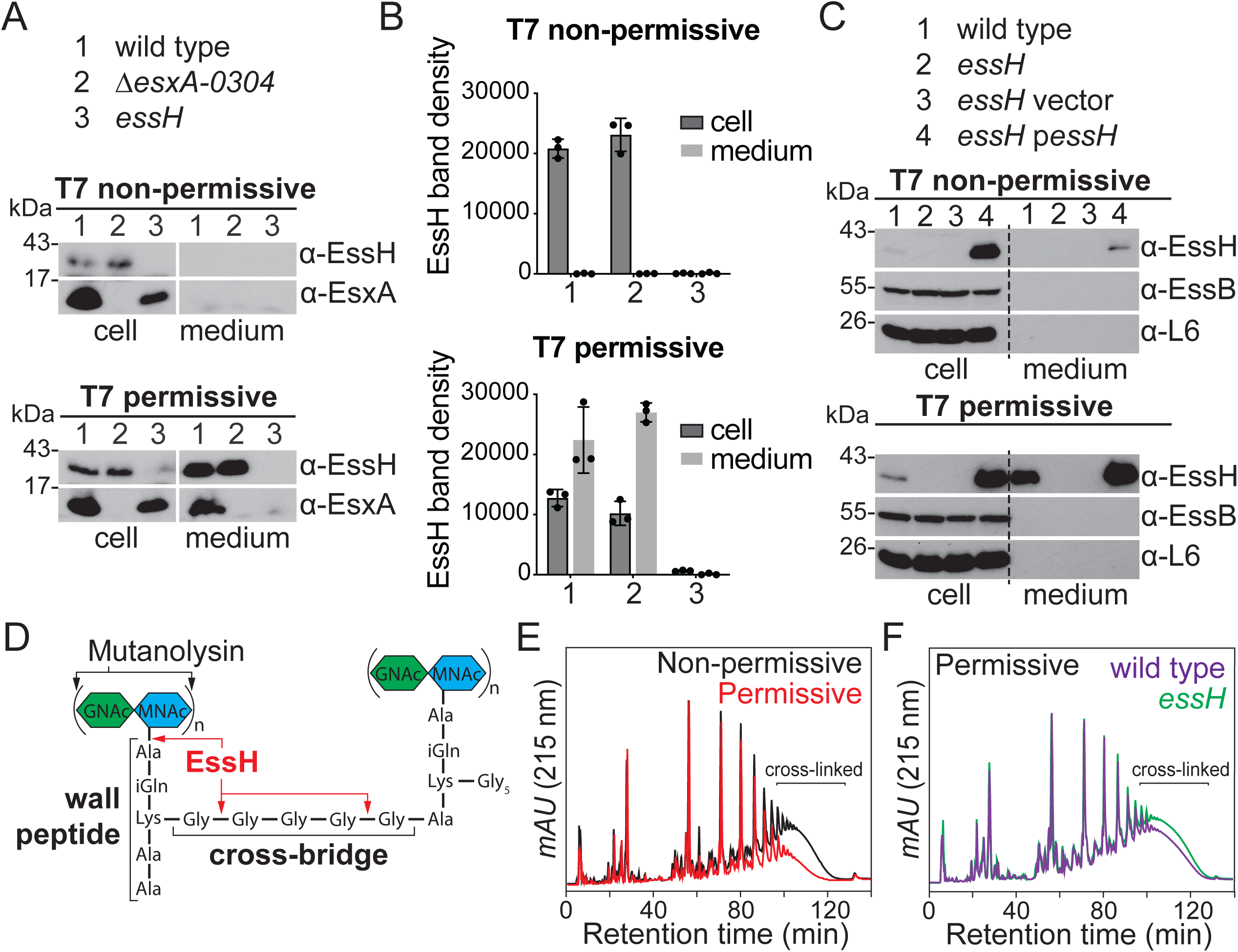
EssH is detected in the culture medium under T7 permissive conditions. **A)** To assess protein secretion, cultures of *S. aureus* USA300 LAC* (wild type) or its isogenic Δ*esxA*-*0304* and *essH* mutants, were cultured in either TSB (T7 non-permissive) or TSB supplemented with heat-inactivated horse serum (HS/HI) at pH5.5 (T7 permissive) to induce T7b secretion. Proteins were prepared from lysed cells and medium fractions, resolved by SDS-PAGE and analyzed by immunoblotting using rabbit polyclonal antibodies specific for EsxA (α-EsxA) and EssH (α-EssH). Numbers (in kDa) indicate the migratory positions of molecular weight markers on SDS-PAGE. **B)** EssH band densities from three independent experiments in (A) were quantified and plotted for cultures growing under T7 non-permissive and T7 permissive conditions. **C)** Wild type, *essH*, and *essH* bearing pSEW016 (vector) or p*essH* plasmids were cultured, fractionated and analyzed as described above. Polyclonal rabbit antibodies specific for EssH (α-EssH), EssB (α-EssB) and ribosomal protein L6 (α-L6) were used for immunoblot analyses. **D)** Diagram of cross-linked *S. aureus* peptidoglycan with black arrows specifying the bonds cleaved by mutanolysin and red arrows specifying the bonds cleaved by EssH. Green and blue hexagons represent *N*-acetylglucosamine (GlcNAc) and *N*-acetylmuramic acid (MurNAc), respectively. **E)** Wild type was cultured under T7 non-permissive (black trace) and T7 permissive (red trace) conditions and murein sacculi prepared, glycan chains digested with mutanolysin and separated by C^18^ reversed-phase HPLC to assess the extent of cross-linking. **F)** Wild type and *essH* strains were cultured under T7-permissive conditions, murein sacculi prepared and analyzed like in (E). mAU corresponds to milli- absorbance units measured at 215 nm wavelength.

EssH is a cell wall hydrolase with specific amidase and endopeptidase activities (Fig. 1D), which affects the cross-linking of the adjacent peptidoglycan subunits (37). To test whether the difference in EssH secretion under the two conditions affects the cross-linking of the PG *in vivo*, we analyzed the composition of murein sacculi from cells cultured under T7 permissive and non-permissive conditions. Murein sacculi were purified from *S. aureus* and treated with mutanolysin which cleaves β-N-acetylmuramyl-(1→4)-N-acetylglucosamine linkages within the glycan chains (Fig. 1D), allowing to resolve and assess crosslinked muropeptides by C^18^ reversed-phase high-performance liquid chromatography (HPLC). PG of staphylococci from T7 permissive conditions is less cross-linked compared to that of cells grown under T7 non-permissive conditions (Fig. 1E). To test whether this marked reduction in PG cross-linking is dependent on EssH, we analyzed murein sacculi from wild type and *essH* mutant strains grown under T7 permissive conditions. The *essH* mutant had a modest increase in PG cross-linking compared to wild type, indicating that EssH is partially responsible for the observed changes in PG cross-linking under T7 permissive conditions (Fig. 1F). Taken together, these results reveal that EssH secretion and its cell wall hydrolytic activity increase under T7 permissive growth conditions.

### Stability of EssH is controlled by *S. aureus* secreted protease

To better understand how T7 permissive conditions impact EssH secretion, culture media were adjusted either for pH or HS/HI. Immune detection of EssH in culture media was dramatically increased following growth of bacteria in the presence of HS/HI while lowering the pH only had a minor effect (Fig. 2A). Since EssH is secreted via the Sec pathway, it was not immediately clear how HS/HI would selectively affect its secretion. We reasoned that secreted EssH may simply be unstable and rapidly degraded by proteases also secreted in the culture medium. Indeed, horse serum is rich in albumin and globulin (𝛼_1_, 𝛼_2_, 𝛽_1_, 𝛽_2_, 𝛾) proteins (38) that could act as competing substrates for proteolysis. To test this hypothesis, wild type bacteria were grown in TSB supplemented with a protease inhibitor (PI) cocktail (Fig. 2A). An EssH immune signal was indeed observed, suggesting that EssH may be susceptible to proteolysis.

**Figure 2.**
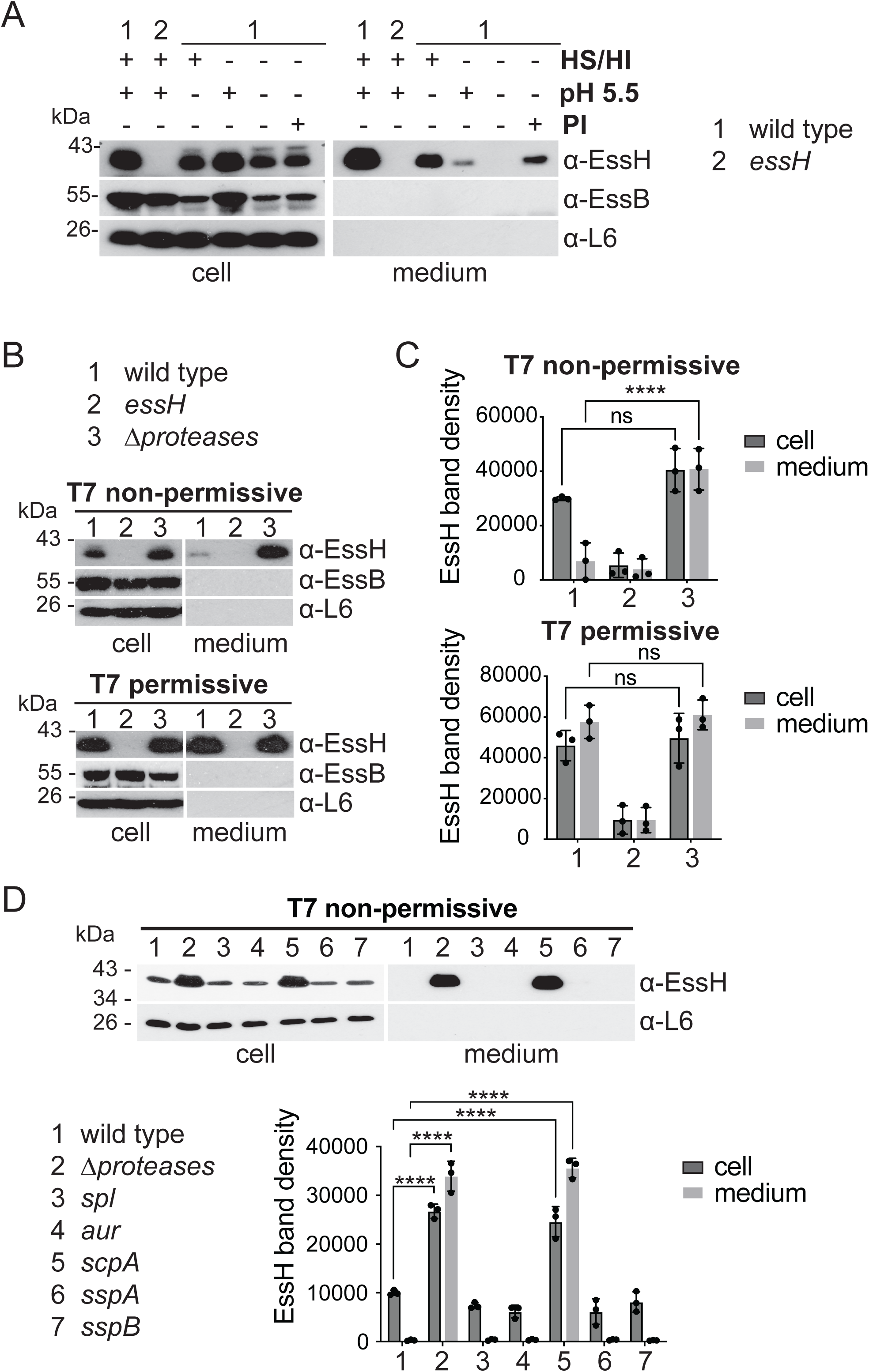
EssH is stabilized in *scpA::ermB* mutant under T7 non-permissive conditions. **A)** *S. aureus* wild type or *essH* control were cultured in TSB medium, or TSB supplemented with HS/HI at pH5.5, HS/HI only, pH5.5 only, or protease inhibitors (PI). Cultures were fractionated and proteins analyzed by immunoblot using polyclonal rabbit antibodies specific for indicated proteins. **B)** Wild type, *essH* and Δ*proteases* were cultured under T7 non-permissive or T7 permissive conditions. Cultures were fractionated and proteins in cell and medium analyzed by immunoblot using polyclonal rabbit antibodies raised against indicated proteins. **C)** EssH band densities from triplicate experiments in (B) were quantified and plotted. Data was analyzed for statistical significance using two-way ANOVA followed by Tukey’s multiple comparison test; p-values 0.1234 (ns), <0.0001 (****). **D) W**ild type and isogenic mutants Δ*proteases* (AH1919), *spl* (DM2288), *aur* (BDG2485), *scpA* (BDG2486), *sspA* (BDG2487) and *sspB* (BDG2488) were cultured under T7 non-permissive conditions, fractionated and analyzed by immunoblot using polyclonal rabbit antibodies specific for indicated proteins. EssH band densities from triplicate experiments were quantified and plotted. Data was analyzed for statistical significance by two-way ANOVA followed by Tukey’s multiple comparison test; p-values <0.0001 (****).

*S. aureus* USA300 LAC* secretes ten major proteases, SplA-F (serine protease-like proteins A to F), Aur (aureolysin), ScpA (staphopain A), SspA and SspB (serine proteases A and B), that have been shown to degrade both staphylococcal and host proteins (39). The Δ*proteases* mutant lacks all ten secreted proteases and was used to compare the abundance of EssH in media of T7 non-permissive versus T7 permissive cultures. This analysis clearly demonstrated that the inability to observe EssH by western blot in our so-called T7 non-permissive conditions correlates with the simultaneous secretion of proteases (Figs. 2B and 2C). To examine if one of the ten proteases may be responsible for the degradation of EssH, we probed *S. aureus* mutants with transposon (Tn::*ermB*) insertions disrupting genes *aur*, *scpA*, *sspA* and *sspB,* or lacking the *splABCDEF* operon. EssH could be clearly detected in the supernatant of the mutant lacking *scpA* grown under T7 non-permissive conditions, in fact the relative abundance of EssH also increased in the cell fractions of both *scpA* and Δ*proteases* mutants (Fig. 2D). Taken together, these findings indicate that secreted EssH is unstable and degraded by ScpA, decreasing the pH may reduce the activity of ScpA and growing bacteria in the presence of serum provides competing substrates.

### Free EssH found in the medium is expendable for T7b-dependent secretion

In Fig. 1A, we show that EssH is required for the secretion of EsxA and T7 secretion correlates with the presence of EssH in the culture medium under T7 permissive growth. We wondered if the accumulation of EssH in Δ*proteases* and *scpA* mutants is sufficient to support EsxA secretion in otherwise T7 non-permissive conditions. However, this was not the case. EsxA was not detected in the medium fractions of the Δ*proteases* and *scpA* mutants despite the presence of EssH (Fig. 3A; medium lanes 4 and 5). Fractionations of strains Δ*esxA-0304* and Δ*essH* are shown as controls (Fig. 3A; medium, lanes 2 and 3). These results indicate that the removal of proteases to stabilize secreted EssH is not sufficient to support T7b secretion under T7 permissive conditions.

**Figure 3.**
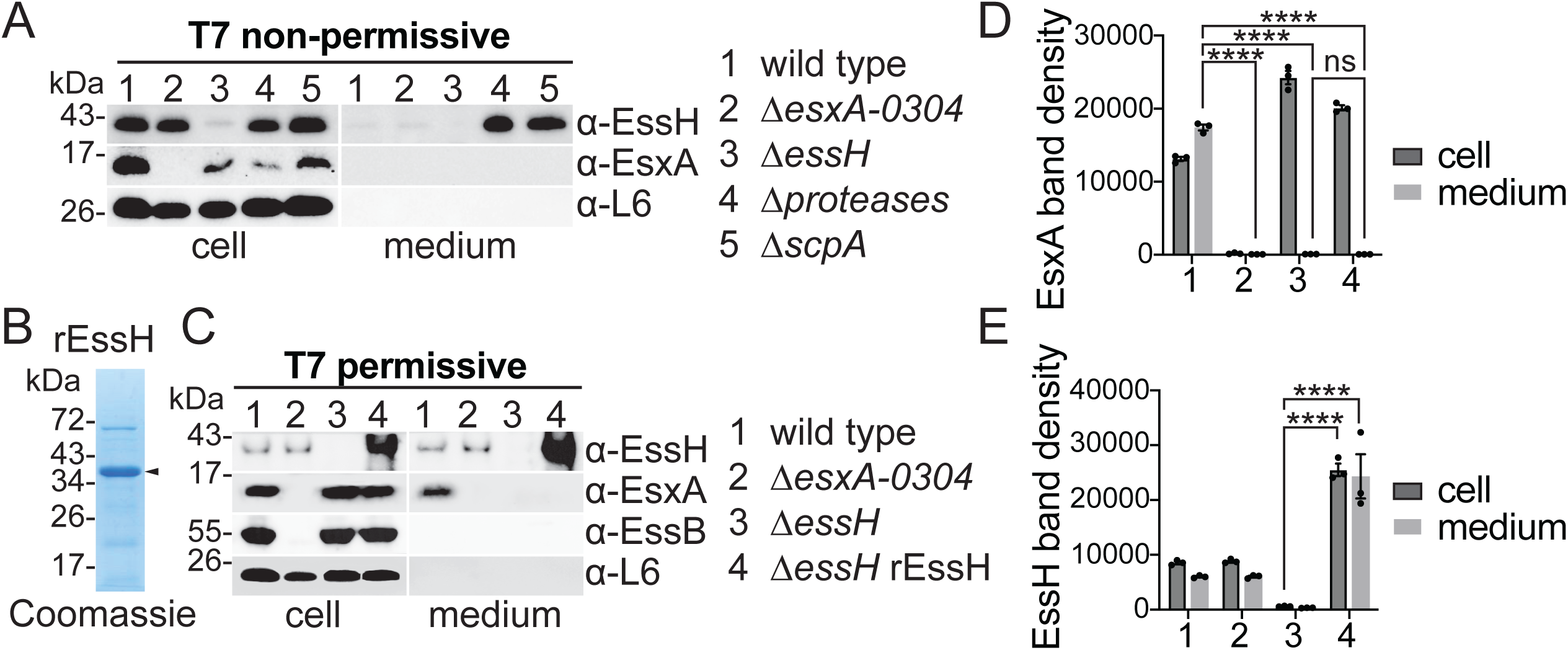
Conditional induction of T7b-dependent secretion does not rely on staphylococcal secreted proteases and free EssH in the mileu. **A)** Wild type, Δ*essB, ΔessH*, Δ*proteases* and Δ*scpA* were cultured under T7 non-permissive conditions and fractionated into cell and medium. Proteins in each fraction were analyzed by immunoblot using polyclonal rabbit antibodies specific for indicated proteins. **B)** Recombinant EssH (rEssH) protein was purified from *E. coli* BL21 p_STREP_*essH* strain as previously described (37), resolved by SDS-PAGE and stained with Coomassie. Black arrow indicates rEssH. C) *S. aureus ΔessH* was grown under T7 permissive conditions in the absence (lane 3) or presence (lane 4) of the purified rEssH protein added in the culture medium. As secretion controls, wild type and Δ*esxA-0304* were cultured under T7 permissive conditions without rEssH supplementation. Cultures were fractionated into cell and medium. Proteins were prepared from each fraction and analyzed by immunoblot using polyclonal rabbit antibodies specific for indicated proteins. **D)** EsxA and **E)** EssH band densities from three independent experiments in (C) were quantified and plotted. Data was analyzed for statistical significance by two-way ANOVA followed by Tukey’s multiple comparison test; p-values 0.1234 (ns), <0.0001 (****).

Next, we asked if adding mature, recombinant EssH (rEssH) purified from *E. coli* (Fig. 3B) to the culture of the Δ*essH* mutant grown under T7 permissive conditions, would restore EsxA secretion (Fig. 3C-E). This was also not the case, supplementing the culture in this manner did not restore EsxA secretion as shown by western blotting and by quantifying immune signals (Fig. 3C-E). To ascertain that purified rEssH was enzymatically active, we performed an *in vitro* assay. *S. aureus* murein sacculi were purified, treated with mutanolysin and then incubated with rEssH or buffer before analysis by C^18^ reversed-phase HPLC. PG treated with rEssH had reduced cross-linking and produced earlier eluting muropeptides (Fig. S1) which are consistent with the amidase and endopeptidase activities of the EssH CHAP domain that were characterized in our prior study (37). Together these results indicate that while under T7 permissive conditions or in absence of proteases EssH is quite abundant in the medium, this free secreted form may no longer contribute to T7 secretion. Thus, we surmise that any key contribution of EssH for T7b secretion must occur prior to its release into the extracellular milieu.

### EssH associates with the T7SSb membrane complex

If adding rEssH to cultures does not result in cell lysis and does not rescue T7b secretion in cells lacking *essH*, we must infer that the PG hydrolase activity of EssH acts while the protein is still in transit in the murein sacculus. Another reasonable assumption is that secreted EssH also interacts with the T7b secretion machinery. Previously, we expressed EssC with a C-terminal Twin-Strep tag, EssC_TS_, in *S. aureus* and copurified the membrane-spanning subunits EsaA, EssA, EssB and EssC_TS_ over Strep-tactin Sepharose (20, 21). Here, plasmid-encoded *essC*_TS_ or plasmid-encoded *essC* without a tag (serving as a control) were transformed in wild type and the Δ*essH* mutant. Affinity chromatography over Strep-tactin Sepharose was used to purify EssC or EssC_TS_ from detergent-solubilized membranes of wild type and Δ*essH* mutant. Bound proteins were eluted, separated by SDS-PAGE and transferred for immunoblot analysis using specific rabbit polyclonal antibodies. As expected, EssC_TS_ co-purified with the components of the T7SSb membrane complex, namely EsxA, EsaA, EssA and EssB, in both wild type and Δ*essH* mutant (Fig. 4A). Importantly, EssH was also found associated with this complex purified from wild type extracts but was absent from Δ*essH* mutant extracts as well as the tag-less EssC control (Fig. 4A). These results indicate that EssH associates with the T7SSb complex in the cell envelope.

**Figure 4.**
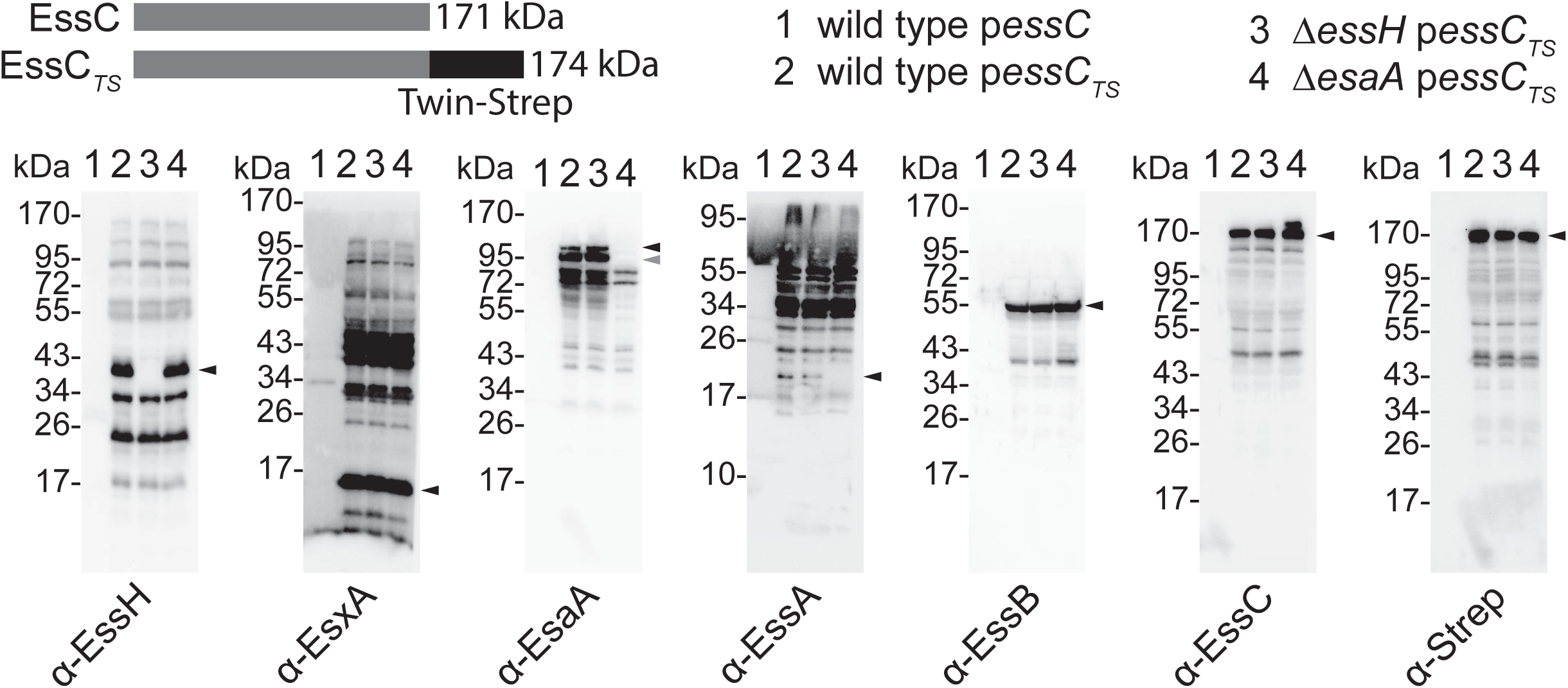
EssH co-purifies with the T7SSb complex. EssC or EssC_TS_ variant bearing a C-terminal Twin-Strep-tag (top left) were used to purify T7SSb complex. Wild type carrying p*essC* (control) or wild type, Δ*essH* and Δ*esaA* carrying p*essC_TS_* were cultured under T7 permissive conditions. Detergent extracts from staphylococcal membranes were prepared and purified by affinity chromatography over Strep-Tactin Sepharose resin. Proteins in eluates were separated by SDS-PAGE, electrotransferred to PVDF membranes for immunoblot analyses with indicated polyclonal antibodies. Black arrows indicate immunoreactive species corresponding to the expected size of the proteins of interest. Grey arrow indicates an alternative immunoreactive EsaA species. Numbers (in kDa) indicate the migratory positions of molecular weight markers. Experiment was performed in triplicate and representative western blots are shown.

### EssH is required for EsaA secretion

EsaA contains 6 TM domains and a long soluble domain that connects TM1 and TM2. Western blotting previously revealed that EsaA is processed into two stable products that fractionate to the cell and medium of wild type bacterial cultures. Because EsaA is a membrane protein, we reasoned that its shedding into the milieu may represent an end product of the T7SSb. This shedding also suggests that most of cell-associated EsaA is embedded in the cell wall. While we do not fully understand the mechanism of processing, we wondered if EssH could contribute to shedding into the milieu. In line with this prediction, the abundance of secreted EsaA species was indeed diminished in the medium but unchanged in the cells of the Δ*essH* mutant when compared to wild type (Figs. 5A and 5B). As before, secretion of EsxA was also inhibited in Δ*essH* mutant cells. Complementation of Δ*essH* with the p*essH* plasmid restored the release of EsaA species in the milieu (Figs. 5C and 5D). Thus, release of EsaA from the cells is no longer observed in a mutant lacking *essH*, but EsaA processing is independent of EssH and of the secreted proteases Aur, ScpA, SspA, SspB, SplA-SplF (Figs. 5A and 5B).

**Figure 5.**
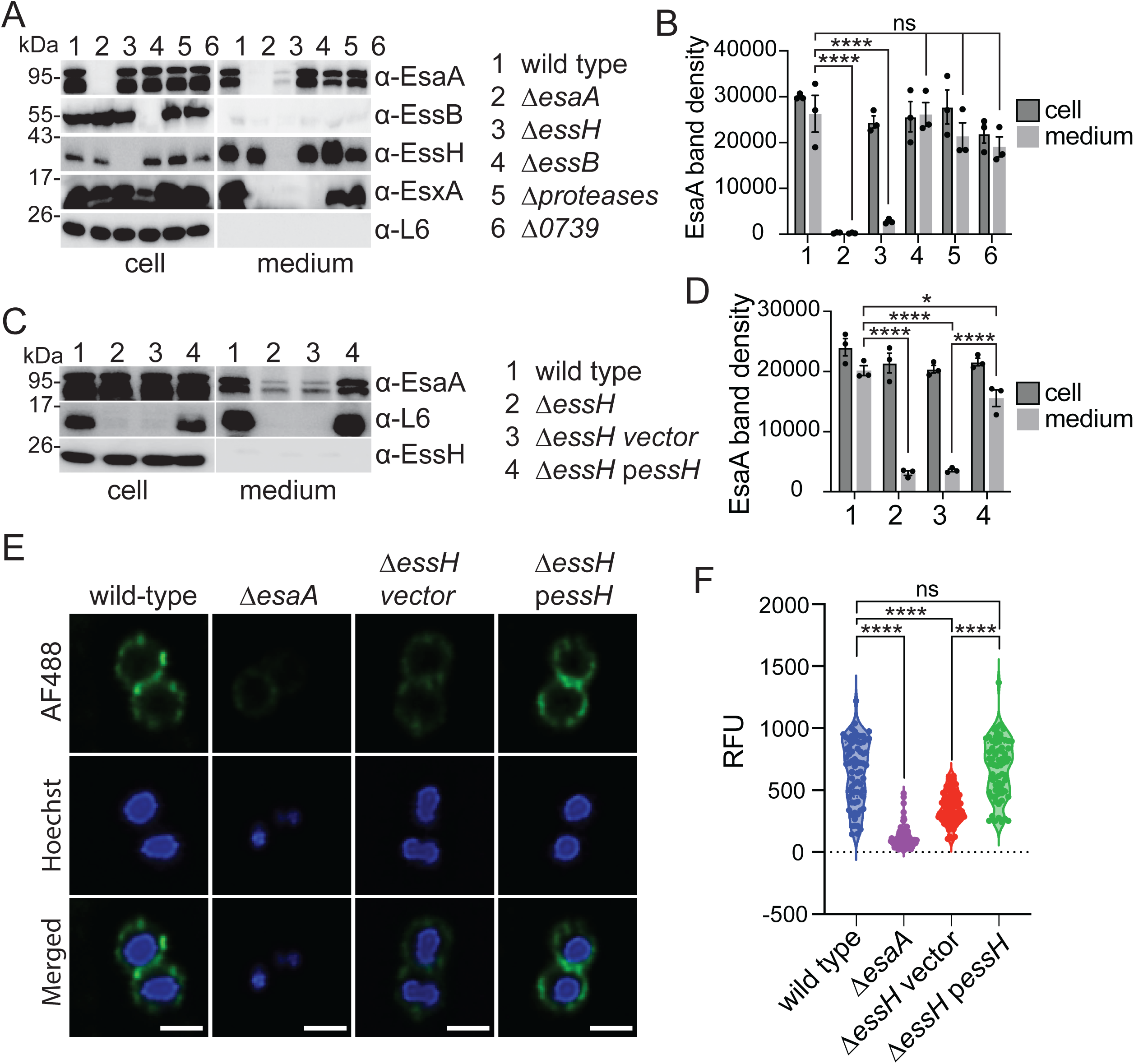
EssH is required for EsaA secretion and surface exposure. **A)** Wild type, Δ*esaA,* Δ*essH*, Δ*essB*, Δ*proteases* and Δ0739 strains were cultured under T7 permissive conditions and fractionated into cell and medium. Medium fraction was concentrated 25x and both fractions analyzed by immunoblot using specified polyclonal rabbit antibodies. Numbers (in kDa) indicate the migratory positions of molecular weight markers. **B)** Quantification of the EsaA band densities in cell and medium fractions from experiment in (A) performed in triplicate. Data was analyzed for statistical significance by two-way ANOVA followed by Tukey’s multiple comparison test; p-values 0.1234 (ns), <0.0001 (****). **C)** Wild type, Δ*essH,* Δ*essH vector and ΔessH pessH* cultures grown under T7 permissive conditions were fractionated and analyzed same as in (A). **D)** EsaA band densities in cell and medium fractions from triplicate experiments in (C) were quantified and statistical significance determined using two-way ANOVA followed by Tukey’s multiple comparison test; p-values 0.1234 (ns), <0.0332 (*), <0.0001 (****). **E)** Wild type, Δ*esaA*, Δ*essH* vector and Δ*essH* p*essH* strains were washed and fixed. To immunostain, cells were first blocked with 3% BSA and monoclonal human IgG, then incubated with polyclonal rabbit serum specific for EsaA followed by washing and incubation with anti-rabbit monoclonal antibody conjugated to Alexa Fluor 488 (AF488). DNA was stained with Hoechst DNA dye and staphylococci were imaged using Nikon Eclipse Ti2 scanning confocal microscope and representative images shown. Scale bar represents 1 µm. **F)** Fluorescence (RFUs) of cells (n=70) from micrographs collected in three independent experiments in (A) was measured using Fiji software and plotted. Statistical significance was determined by two-way ANOVA with Tukey’s multiple comparison test; p-values 0.1234 (ns), <0.0001 (****).

Next, we use immunofluorescence microscopy to examine how EssH may impact the association of EsaA with cells, using a polyclonal serum specific for the soluble domain of EsaA (α-EsaA) and a secondary anti-rabbit antibody conjugated to Alexa Fluor 488 (AF488). Hoechst fluorescent dye was used as a control to dye DNA. EsaA-AF488-fluorescence was mostly seen surrounding wild type cells, while fluorescence signals were not significant for Δ*esaA* cells and significantly reduced for Δ*essH* cells (Figs. S2A and S2B; Figs. 5E and 5F). EsaA-AF488-fluorescence was restored to near wild type level in the complemented strain Δ*essH* p*essH* (Figs. 5E and 5F). We wondered if EssH might directly interact with the EsaA subunit of T7b machinery. To test this possibility, plasmid-encoded *essC*_TS_ was transformed in the Δ*esaA* mutant and proteins in detergent-solubilized membranes purified over Strep-tactin Sepharose. This approach revealed that EssH is still pulldown with EssC_TS_ along with EssB, and EsxA even in the absence of EsaA; of note and as was shown previously (21) EssA was also not detected in this complex (Fig. 4A; lanes 4). Taken together, we conclude that a large portion of EsaA spans the cell wall envelope and while EssH does not directly associate with the EsaA, it contributes to the envelope localization of EsaA.

### The N-terminal Domain of EssH (EssH ND) is necessary to support T7b-dependent secretion

The C-terminal segment of EssH amino acids 151 through 297 is a canonical CHAP domain with the catalytic residues, Cys^199^ and His^254^, responsible for the PG hydrolytic activity of EssH (37). The first 24 N-terminal amino acids of EssH comprise a signal peptide for Sec-dependent secretion. Amino acids 25 to 150 do not share any conserved features and this domain is referred to as the N-terminal domain (ND). To test the requirement of the ND and CHAP domains for T7SSb, the Δ*essH* mutant was complemented with plasmids carrying either full-length *essH* (p*essH*), or variants deleted for the ND (p*essH^ΔND^*) or with mutations in the catalytic residues C199A and H254A (p*essH^C199A/H254A^*) (Fig. 6A). Only the plasmid encoding full length EssH complemented the T7b secretion defect of the Δ*essH* (Figs. 6B-D). Similarly, strains e*ssH^ΔND^* and *essH^ΔCHAP^* that produce genome encoded EssH lacking the ND or CHAP domains, respectively, did not secrete EsxA and EsxC (Fig. S3B). But the secretion defects in these strains could be restored by plasmid-encoded EssH suggesting that the truncated proteins EssH^ΔND^ and EssH^ΔCHAP^ did not exert any dominant negative effect (Fig. S3B). To eliminate the possibility that the deletion of ND negated the PG hydrolytic activity of the CHAP domain encoded by p*essH^ΔND^*, we tested whether mutant proteins lacking either ND or CHAP domain could hydrolyze staphylococcal PG. Affinity chromatography over Strep-tactin Sepharose was used to purify rEssH, rEssH^ΔND^ and rEssH^ΔCHAP^ (Figs. 6E and 6F). Purified proteins were then incubated with mutanolysin-treated peptidoglycan to test their hydrolytic activity (Fig. 6G). Both, rEssH and rEssH^ΔND^ hydrolyzed PG, producing muropeptide fragments consistent with amidase and endopeptidase activity of the CHAP domain as determined previously (37). Deletion of the CHAP domain in rEssH^ΔCHAP^ abrogated its hydrolytic activity producing absorbance profile similar to that of mutanolysin-treated control (Fig. 6G). Taken together, EssH CHAP hydrolytic activity is necessary but not sufficient to support T7b secretion and requires the sequence contained within the ND. To further query the specific requirement of the CHAP domain of EssH, we examined the secretion of EsaA and EsxA in the Δ*0739* mutant (Figs. 5B and 5C). Gene *SAUSA300_0739* encodes an uncharacterized but conserved PG hydrolase with a conserved CHAP domain. We observed that *SAUSA300_*0739 is dispensable for the secretion of EsaA and EsxA (Figs. 5B and 5C), highlighting the T7SSb specificity of EssH.

**Figure 6.**
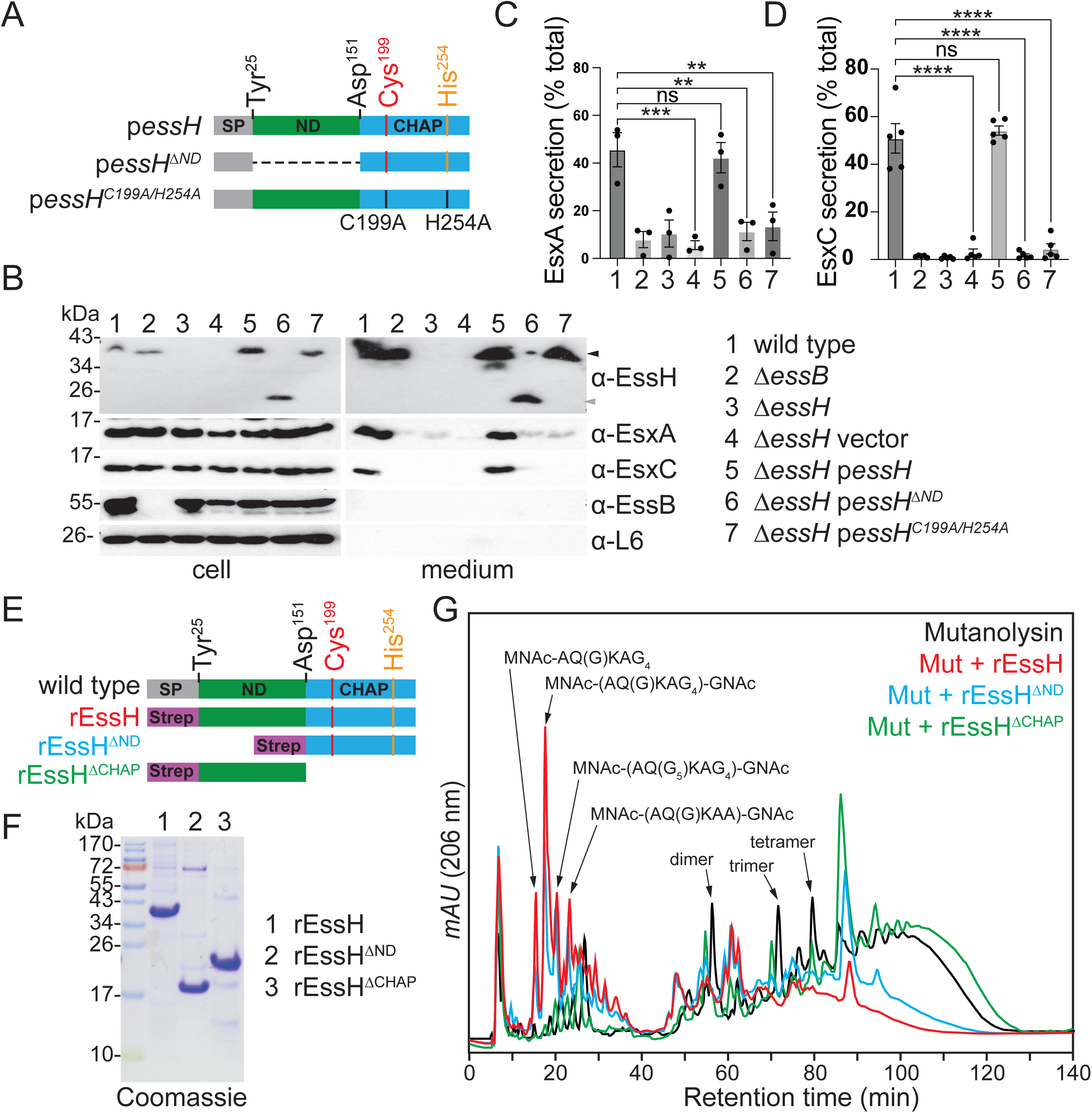
EssH N-terminal domain (ND) is required for T7b-dependent secretion. **A)** Illustration of EssH variants encoded by plasmids p*essH*, p*essH^ΔND^* and p*essH^C199A/H254A^* with annotated SP, ND and CHAP domains, as well as catalytic Cys^199^ and His^254^ residues with C199A and H254A mutations. **B)** Strains wild type, *ΔessB, ΔessH* and *ΔessH* bearing vector, p*essH*, p*essH^ΔND^* and p*essH^C199A/H254A^* plasmids were cultured under T7 permissive conditions, fractionated and analyzed by immunoblot using specified polyclonal rabbit antibodies. **C-D)** Densitometry quantification of the abundance of C) EsxA and D) EsxC immune-reactive signals in cell and medium fractions from three independent experiments in (B). Data are analyzed as percentages of total EsxA or EsxC respectively, i.e. the added densities of protein in cell and medium fractions. Statistical significance was determined by ordinary one-way ANOVA with Dunnett multiple comparisons test; p-value 0.123 (ns), 0.0021(**), 0.0002(***), <0.0001(****). **E)** Diagram of wild type EssH protein and purified variants rEssH, rEssH^ΔND^ and rEssH^ΔCHAP^ appended with a N-terminal Step-tag II. Residues Tyr^25^ and Asp^151^ denote beginning of ND and CHAP domains respectively, and Cys^199^ and His^254^ denote catalytic residues. **F)** Coomassie brilliant blue-stained SDS-PAGE gel separating purified rEssH, rEssH^ΔND^ and rEssH^ΔCHAP^. Numbers (in kDa) indicate the migratory positions of molecular weight markers. **G)** Purified *S. aureus* peptidoglycans were digested with mutanolysin and split into two samples that were either left untreated (black trace) or were treated with purified rEssH (red trace), rEssH^ΔND^ (blue trace) and rEssH^ΔCHAP^ (green trace). Resulting muropeptides were separated using C^18^ reversed-phase HPLC. Expected structures of muropeptides in the corresponding peaks were denoted based on the prior analysis of mutanolysin and rEssH treated peptidoglycan (37). mAU corresponds to milli-absorbance units measured at 206 nm wavelength.

## DISCUSSION

In *S. aureus*, T7SSb assembles in the cell envelope into a core complex comprised of EsaA, EssA, EssB and EssC proteins (20, 21). The staphylococcal cell wall is 40-50 nm thick, with an estimated pore size of ∼2 nm, which might preclude the assembly of the T7SSb complex or trafficking of secreted proteins (40, 41). Nonetheless, T7SSb is predicted to extend past the peptidoglycan via its EsaA subunit. EsaA is not conserved in the T7SSa of actinobacteria and contains six TM domains (TM1-6), with a 725 amino acid-long soluble domain separating TM1 and TM2. The soluble domain of *S. gallolyticus* EsaA was shown to form arrow-like dimers approximately ∼20 nm long, which in the context of the full protein are predicted to span the length of the firmicute cell wall (24). In *B. subtilis*, the EsaA homolog YueB serves as a receptor for bacteriophage SPP1, further supporting that EsaA spans the cell wall and is surface-exposed (25). In Gram-negative organisms, molecular complexes that span the periplasm such as type IV pili, as well as type III, IV and V secretion systems are equipped with lytic transglycosylases that locally cleave β-(1→4)-glycosydic bonds within the (GlcNAc-MurNac)_n_ strands of PG (42). For example, the lytic transglycosylase EtgA interacts at 1:1 ratio with EscI, the inner rod component of the type III secretion system (T3SS) needle complex. This interaction triggers EtgA murein hydrolase activity to help T3SS inner rod assemble within the cell wall (43). EssH is unlike EtgA as it is a non-lytic PG hydrolase associated with the ESS gene cluster of *S. aureus* (37). EssH is one of ten conserved secreted PG hydrolases with C-terminal CHAP domains, characterized by the presence of conserved catalytic Cys and His residues (44–46). Deletion of individual CHAP proteins is associated with changes in sensitivity to cell wall-targeting antibiotics, biofilm formation, protein secretion and cell division (47–50). However, *S. aureus essH* mutants specifically fail to secrete T7SSb substrate proteins such as EsxA, EsxC and EssD (EsaD) (37). Minor changes in PG crosslinking in the *essH* mutant suggest that EssH might act locally to remodel PG at the site of T7SSb assembly (Fig. 1F). In agreement with this notion, we find that EssH associates with the T7SSb complex when purified from detergent-soluble membranes by affinity chromatography of EssC_TS_ (Fig. 4). Surprisingly, this association was retained in the Δ*esaA* mutant (Fig. 4), implying that EssH likely interacts with another subunit of the T7SSb complex. While EssH does not appear to interact directly with EsaA, both cell surface display and shedding of EsaA require EssH (Fig. 5A-D). However, EsaA still copurifies with the T7SSb complex in the absence of Δ*essH*, suggesting that EssH functions to extend EsaA across the cell wall but does not interfere with EsaA assembly with the other T7SSb core components within the membrane. Thus, EssH is secreted via the SecYEG translocon and associates with the T7b machinery. We speculate that this association allows local PG hydrolysis and assembly of the T7SSb complex across the cell wall, specifically its EsaA subunit (Fig. 7A); in the absence of EssH, the secretion of substrates is abrogated because EsaA is unable to extend the T7b conduit across the cell wall (Fig. 7B).

**Figure 7.**
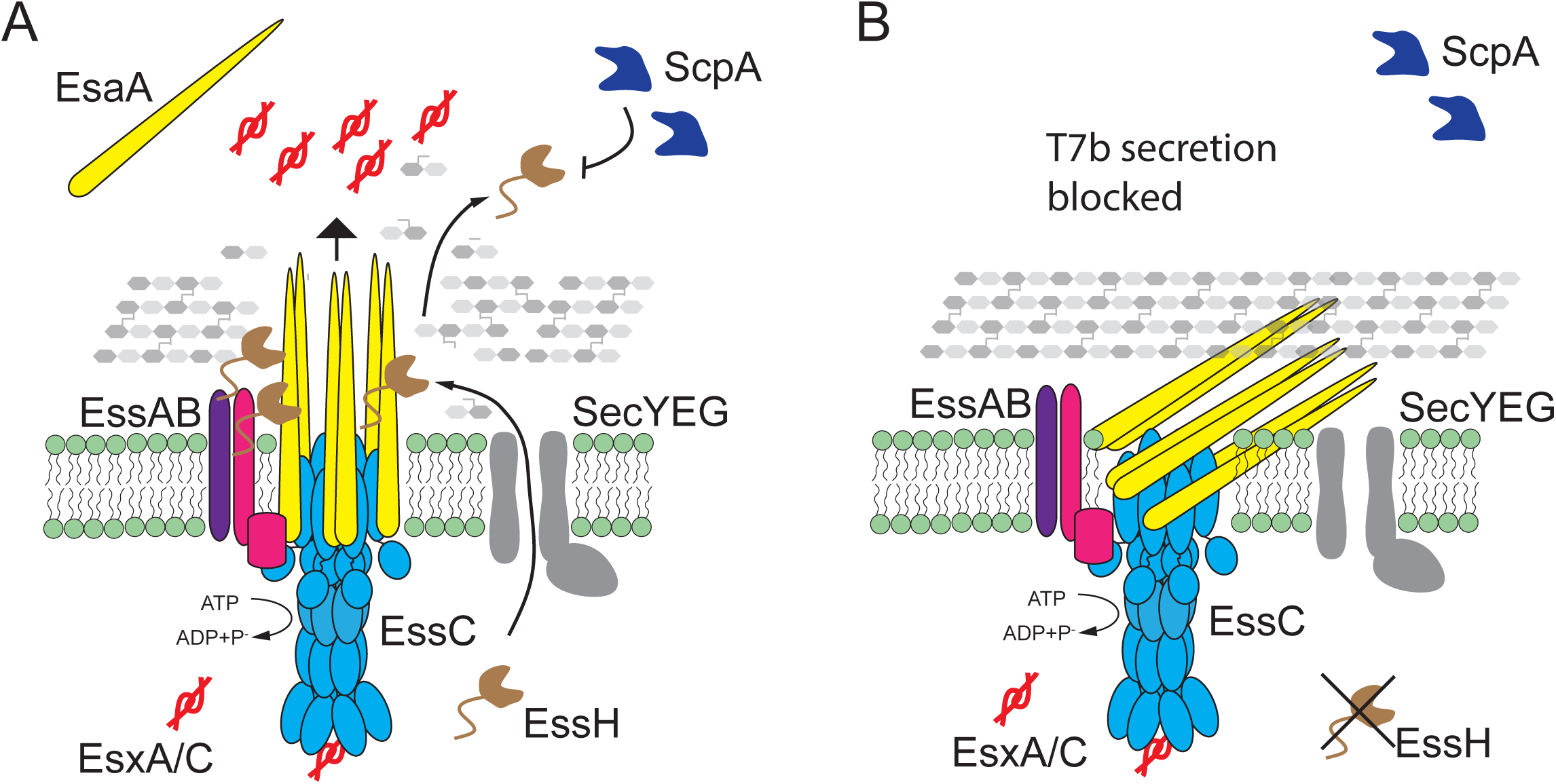
Model of EssH function to support secretion via T7SSb. **A)** An illustration of T7SSb complex consisting of EsaA (yellow), EssA (purple), EssB (pink) and EssC (light blue) membrane proteins that assemble across *S. aureus* cell envelope to support secretion of EsxA and EsxC (EsxA/C;red) substrates. EssH (brown) is translocated across the membrane via general secretion pathway translocase (SecYEG; grey). EssH interacts with the T7SSb complex and hydrolyzes the PG to allow for EsaA to assemble across the cell wall. EssH is then released into the milieu where it becomes susceptible to proteolysis by the secreted cystine protease ScpA (Staphopain A; dark blue). **B)** In the absence of EssH, thick PG layer prevents accommodation of a large EsaA subunit, which prevents EsaA shedding into the milieu and blocks secretion of EsxA/C substrates.

It is not yet clear how EssH associates with the T7SSb. The predicted structure of EssH generated by AlphaFold (51) reveals that ND is unstructured (Fig. S4). Deletion of ND (*essH^ΔND^*) abrogates EsxA and EsxC secretion even though the isolated CHAP domain can still hydrolyze PG (Figs. 6B, 6G, S3B and S3C). Thus, both domains, PG hydrolytic CHAP and unstructured ND, are required for T7b-mediated secretion by EssH. Unstructured regions within proteins referred to as intrinsically disordered regions (IDRs) are found in all domains of life (52). IDRs often contain functional modules that mediate protein-protein interactions, such as short linear motifs (SLiMs) and intrinsically disordered domains (IDDs) (53, 54). These modules can determine protein stability, serve as sites for post-translational modifications, alter protein function, determine protein localization, enable concentration-dependent phase transitions or mediate dynamic interactions within protein complexes (55, 56). In bacteria, the functions of only a few IDRs are known, limiting our understanding of their physiological implications. For example, in *B. subtilis* IDRs are involved in cell wall homeostasis and response to stress. Cell wall stress in the form of gaps in the PG meshwork is sensed via regulated intramembrane proteolysis of RsgI, an anti-σ factor, which then derepresses the stress response σ^I^. It has been proposed that the IDR of RsgI extends into the cell wall and detects PG defects, which leads to the regulated proteolytic cleavage of RsgI and the activatation of the stress response (57). In *Bacteroides thetaiotaomicron,* the Rho protein responsible for Rho-dependent termination of transcription contains an IDR which enables formation of cytosolic droplets or compartments through liquid-liquid phase separation. Deletion of this IDR results in the loss of droplet formation by Rho and mutant bacteria have reduced fitness in the mouse model of gut colonization (58). Analysis of the EssH sequence using FuzDrop (59) identified a droplet promoting region spanning amino acids 59 to 124 within the ND domain, indicating it has potential for liquid-liquid phase separation. While the significance of this motif is not known, it could help increase the concentration of EssH near the T7b complex providing a mean to locally remodel the PG. Alternatively, ND could monitor the interaction of EssH with the T7b complex to modulate the activity of the CHAP domain.

The model where EssH locally hydrolyzes PG at the site of T7SSb assembly also explains why it becomes dispensable upon secretion into the milieu (Figs. 2B and 2C). Secreted EssH released into the milieu is rapidly degraded by secreted proteases (Fig. 2B). *S. aureus* encodes ten conserved papain-like secreted proteases, aureolysin (Aur), serine V8 protease (SspA), serine-like proteases (SplA-SplF), and cysteine proteases SspB and Staphopain A (ScpA) (39). Deletion of all ten proteases (Δ*proteases*) leads to a hypervirulent phenotype during septicemia in mice, which has been attributed to the loss of targeted proteolysis of several virulence factors (i.e. SpA, Sbi, IsaB) in this strain (60, 61). Here we found that secreted EssH is stabilized only in the *scpA* mutant, indicating a specific proteolytic targeting by Staphopain A (Fig. 2D). Corroborating our findings, EssH was more abundant in the secretome of a *scpA* mutant strain, whereas it was unchanged in the *aur*, *sspA* and *sspB* mutants (61). Proteases are secreted as pro-enzymes or zymogens that are hierarchically activated by proteolysis. The Aur zymogen is activated by autocatalysis, and then mature Aur activates SspA by limited proteolysis, which in turn activates SspB (62–64). ScpA on the other hand is excluded from this proteolytic cascade and is thought to undergo autocatalysis (65). This would allow ScpA to target EssH independently of the other proteases. Interestingly enough, ScpA-dependent proteolysis of EssH is inhibited by the addition of HS/HI to the culture medium (Fig. 2A). Here, we reasoned that horse serum could provide competing substrates for proteolysis since it contains 51-72 mg/ml of albumin and globulin (𝛼_1_, 𝛼_2_, 𝛽_1_, 𝛽_2_, 𝛾) proteins (38), thereby titrating ScpA away from EssH. Horse serum also contains *cis*-unsaturated fatty acids, such as linoleic acid, that are phosphorylated by fatty acid kinase (Fak) and incorporated into *S. aureus* membrane (66). Linoleic acid was shown to stimulate EsxA secretion in a Fak-dependent fashion, suggesting that membrane incorporation of host unsaturated fatty acids leads to T7SSb activation (67). While the mechanism of HS/HI- dependent activation of T7SSb is not fully understood, it likely does not function via stabilization of secreted EssH, since EsxA was not secreted in the absence of HS/HI independently of EssH abundance in the milieu (Fig. 3A). Moreover, supplementing purified rEssH to the culture medium under T7 permissive conditions failed to complement Δ*essH* mutant and did not restore EsxA secretion (Fig. 3B). Thus, stands to reason that EssH must exert its PG hydrolytic activity to support T7SSb secretion prior to its release into the milieu, at which point it is no longer needed and is degraded by ScpA.

In conclusion, EssH is a conserved PG hydrolase that associates with the T7SSb complex, which allows for EsaA subunit to span the thick cell wall and support secretion of ESS substrates. Both, the ND and CHAP domains of EssH are required to facilitate this process. EssH is then released into the environment where it becomes expendable and is specifically degraded by the secreted protease Staphopain A (ScpA) (Fig. 7A). Future work is needed to characterize the specific interactions of EssH with the T7SSb complex and determine the potential role of the intrinsically disordered ND tail in this process.

## MATERIALS AND METHODS

### Media and growth conditions

*Staphylococcus aureus* was cultured in tryptic soy broth (TSB) or agar (TSA) at 37°C, unless otherwise stated. Media was supplemented with 10 µg/ml chloramphenicol for plasmid selection. Media was adjusted to pH 5.5 and supplemented with 0.2% HS/HI (Gibco) for T7 permissive conditions. *Escherichia coli* was cultured in Lysogeny broth (LB) or agar (LBA) at 37°C, supplemented with 100 µg/ml ampicillin for plasmid selection and 1 mM isopropyl β-D-1-thiogalactopyranoside (IPTG) for production of recombinant proteins.

### Bacterial strains and plasmids

Strains and plasmids are listed in Table S1, and primers are listed in Table S2. *S. aureus* USA300 LAC* (wild type) is a clone of the epidemic community-acquired methicillin-resistant *S. aureus* (CA-MRSA) strain (68) cured of pUSA03 plasmid encoding *ermC* (14), which does not affect T7SSb. Plasmid DNA was passaged through restriction-deficient *S. aureus* RN4220 before transformation into USA300 LAC*. *S. aureus essH* was generated via ϕ85-mediated transduction of transposon disrupted *essH::ermB* allele (37). pKOR1 mediated allelic replacement was performed as previously described (69) and was used to generate Δ*essH*, *essH*^ΔND^ and *essH*^ΔCHAP^ mutants. In short, 1 kb fragments upstream and downstream of the region to be deleted were amplified by PCR from LAC* DNA template using the following primer pairs MBP186F/MBP186R and MBP187F/MBP187R (Δ*essH*), MMP1F/MMP1R2 and MMP2F/MMP2R (*essH*^ΔCHAP^), MBP201F/MKP2R, and MBP199F/MBP199R (*essH*^ΔND^). Amplified fragments for generation of corresponding mutants were fused in a subsequent PCR reaction and cloned into pKOR1 using the BP Clonase II kit (Invitrogen).

Shuttle vectors pWWW412 and its derivative pSEW016 carry P*_hprK_* constitutive promoter and Shine-Dalgarno sequences (70), and were used for complementation studies. In pSEW016, the NdeI cloning site of pWWW412 has been replaced with SacI. Construction of p*essH* and the catalytic mutant p*essH^C199A/H254A^* was described previously (37). To construct p*essH^ΔND^*, signal sequence was PCR amplified from LAC* DNA template with primers MBP37F/MBP61R, and CHAP domain was amplified using primers MBP70F/MBP37R. Fragments were fused by PCR and cloned into pSEW016 using SacI and BamHI restriction sites. For the recombinant production of proteins rEssH^ΔND^ and rEssH^ΔCHAP^ bearing N-terminal Strep-tag II, respective template DNA was amplified by PCR using primer pairs MBP149F/MBP37R and MBP45F/MBP212R, and cloned into pET-15b using NcoI and BamHI restriction sites.

### Fractionation of bacterial cultures and immunoblotting

*S. aureus* cultures were fractionated into medium and cells by centrifugation at 10,000×g for 10 min. Sedimented cells were washed and lysed with lysostaphin (10 µg/ml for 1 h at 37°C). Proteins in in both fractions were precipitated with 10% trichloroacetic acid (TCA), washed in cold acetone, and dried. Precipitates were solubilized in 100µl of 0.5 M Tris-HCl (pH 8.0). Proteins were resolved by SDS-PAGE on 12% or 15% gels and visualized either by staining with Coomassie brilliant blue R-250 or western blot. For western blot, proteins were transferred to polyvinylidene difluoride membrane, blocked for 1 h with 5% dry milk in PBS-T (phosphate buffered saline with 0.1% Tween 20) containing 80 µg/ml of human IgG (Sigma-Aldrich) to block protein A, and incubated with primary polyclonal antibodies at a dilution of 1:5,000 for 1h. Membranes were washed 4 times for 10 min in PBS-T, incubated with 1:10,000 horseradish peroxidase (HRP)-conjugated secondary antibody (Cell Signaling Technology) for 1 h and then washed again 4 times for 10 min in PBS-T. Immunoreactive products were revealed by chemiluminescent detection using SuperSignal West Pico chemiluminescent substrate (Thermo Scientific). The blots were developed on Amersham Hyperfilm ECL (GE Healthcare Life Sciences) or using ChemiDoc imaging system (Bio-Rad).

### T7SSb complex purification

Bacteria were cultured under T7 permissive conditions to OD_600_∼3.0, harvested by centrifugation for 10 minutes at 8,000×g at 4°C and suspended in Buffer A (20 mM Tris pH 8, 300 mM NaCl, 10% v/v glycerol, Pierce EDTA-free protease inhibitors). All the subsequent steps were performed at 4°C. Cells were lysed by bead beating and centrifuged at 10,000×g for 12 minutes. Cleared lysates were centrifugated for 1 hour at 100,000×g at 4°C and supernatants discarded. Remaining pellets were suspended in Buffer A with 0.25% n-dodecyl β-D-maltoside (DDM), incubated on a rotating drum for 1 hour at 4°C and remaining insoluble proteins were removed by centrifugation for 30 minutes at 100,000×g at 4°C. Solubilized proteins were purified by gravity flow affinity chromatography over Strep-Tactin Sepharose. Bound resin was washed with Buffer A containing amphipol A8-35 at 4°C. Proteins were then eluted with Buffer A containing amphipol A8-35 and 5 mM desthiobiotin. Eluted fractions were concentrated by TCA precipitation and probed by western blot as described above.

### Recombinant protein purification

*E. coli* were grown to OD_600_ of 2.0 and centrifuged at 8,000×g for 10 minutes. Sedimented cells were suspended in Buffer A (50 mM Tris-HCl [pH 7.5], 150 mM NaCl), and the resulting suspensions were lysed in a French press at 14,000 lb/in^2^. Unbroken cells were removed by centrifugation at 8,000×g for 15 min, and lysates subjected to ultracentrifugation at 100,000×g for 1 hour at 4°C. Soluble proteins were subjected via gravity flow to chromatography using Strep-Tactin Sepharose (IBA) equilibrated with Buffer A. The columns were washed with Buffer A and eluted with 2.5 mM desthiobiotin (IBA) in Buffer A. Protein concentrations were determined with the bicinchoninic acid assay (Pierce).

### Biochemical assays

*S. aureus* peptidoglycan was prepared as previously described (71), adjusted for concentration by OD_600_ in 100 mM phosphate buffer pH 5.5 and incubated for 18 h at 37°C with 50 units of mutanolysin (Sigma) and reactions quenched at 98°C for 10 min. Samples centrifuged at 15,000×g for 15 min and soluble material was further incubated with rEssH, rEssH^ΔND^, rEssH^ΔCHAP^ or buffer alone as control. All samples were neutralized with sodium hydroxide to reach pH 7.0, dried and reduced via the addition of 250 mM sodium borate and 3–5 mg of sodium borohydride. Samples were incubated for 30 min and reactions stopped by the addition of 20% phosphoric acid to reach pH <4.0 as described (71). Reduced muropeptides were separated by reversed-phase HPLC on a C^18^ column (250 × 4.6-mm ODS-Hypersil; Thermo Scientific) as described previously (72).

### Immunofluorescence microscopy

Bacterial cultures were diluted 1:100 in TSB and allowed to grow with shaking at 37°C to OD_600_∼1.0. Bacteria were pelleted by centrifugation at 10,000 × g for 1 min and washed twice with PBS. Cells were fixed for 20 minutes at room temperature with 500 µL of fixative solution (0.01% glutaraldehyde and 2.5% paraformaldehyde in PBS), washed twice with PBS and resuspended in 150 μL of PBS. Suspension was applied to a poly-L-lysine treated glass slide (MP Biomedicals), washed 3 times with PBS and blocked with blocking buffer (3% BSA in PBS) containing 250 µg/ml monoclonal human IgG for 1 hour. Cells were then incubated for 1 hour with blocking buffer containing appropriate dilution (1:800) of EsaA polyclonal rabbit antiserum, washed 8 times with PBS and incubated in the dark for 1 hour with Alexa Fluor 488-conjugated anti-rabbit antibody (1:500) in blocking buffer. Cells were washed 10 times with PBS and stained with 5 µg/ml Nile red (Sigma) and 50 µg/mL Hoechst 33342 DNA dyes for 5 minutes at room temperature. Afterwards the cells were washed 5 times with PBS and SlowFade Diamond Antifade (Invitrogen) mounting solution was applied to the samples prior to sealing with the glass coverslip. Imaging was done with Nikon Eclipse Ti2 scanning confocal microscope with HC PL APO 63× oil objective (1.4 NA, WD 0.14 mm) utilizing NIS-elements AR image acquisition and analysis software.

### Statistical analyses

Immunoblot densitometry and microscopy immunofluorescence were analyzed using ImageJ and Fiji (73, 74) and graphed and analyzed for statistical significance using Prism (GraphPad Software). Experiments were repeated at least three times and statistical significance calculated as described in the figure legends.

## AKNOWLEDGEMENTS

We are grateful to Maria Krunic and Elena Whitney for technical support. We thank Dr. Lindsey Shaw and Dr. Alexander Horswill for providing protease mutant strains. We thank members of the Missiakas and Bobrovskyy labs for hearty discussion and support. We remember late Olaf Schneewind for his everlasting scientific impact and inspiration.

## FUNDING AND ADDITIONAL INFORMATION

The authors acknowledge support from NIAID, Infectious Diseases Branch R00AI171164 and R01AI038897. The content is solely the responsibility of the authors and does not necessarily represent the official views of the National Institutes of Health.

